# Molecular Subtypes of Anaplastic Gliomas Identified with Somatic-Mutation and Pathway Based Gene Signature

**DOI:** 10.1101/313924

**Authors:** Zabi Wardak, Sunho Park, Hong Zhu, Kevin S. Choe, Tae Hyun Hwang

## Abstract

**Purpose:** Anaplastic gliomas constitute heterogeneous population with variable outcomes and no consensus on therapeutic approach. Molecular profiling may provide prognostication beyond clinical and pathologic factors and help guide treatment decisions.

**Experimental Design:** The Cancer Genome Atlas (TCGA) was utilized to derive a 39-gene low grade glioma-specific gene signature. Consensus clustering based on expression of the signature identified subgroups for 176 patients with anaplastic glioma from TCGA. Overall survival (OS) was analyzed for each subgroup. A total of 68 patients from Repository for Molecular Brain Neoplasia Data (REMBRANDT) were used as an independent validation dataset.

**Results:** Consensus clustering separated the TCGA group into two distinct cohorts. The OS was significantly different between two subgroups, 20 vs. 67 months (p<0.001). On univariate analysis, the molecular subgroup, age, KPS, *IDH1/2* mutation, 1p19q-co-deletion, chemotherapy, and use of both chemotherapy and radiation-therapy were significantly associated with OS. On multivariable analysis, the molecular subgroup remained significant with HR of 2.6 (p=.047, 95%CI [1.01-6.68]). In an independent validation with REMBRANDT, consensus clustering based on the signature successfully identified similarly poor prognostic subgroup with median survival of 14 months and concordance of expression patterns in 21 of the genes.

**Conclusion:** Expression patterns of the 39 gene stratified anaplastic gliomas into two distinct subgroups with substantially different OS. This molecular prognostication was validated in an external dataset. Utilization of molecular subgroup, in addition to known prognostic factors may help define those requiring aggressive therapeutic intervention. Characteristic genes within the poor prognostic group may represent potential targets for therapeutic intensification.

*<u>Source code and dataset used in this work is available for reviewers at:</u>* https://www.taehyunlab.org/ntripath

**Importance of the study:** Despite the revolution of tailored therapy, anaplastic gliomas represent a category of tumors without clear treatment recommendations. While current prognostic factors help guide therapy recommendations, further refinement with the addition of molecular markers can help physicians with treatment recommendations. In this study, we developed a 39 gene prognostic gene signature by utilizing Pan-Cancer TCGA mutation profiles of over 5,000 patients across 19 different TCGA cancer types to stratify patients with anaplastic gliomas. We performed consensus clustering based on gene expression profiles of our 39-gene signature without any consideration of clinical factors or outcomes to TCGA anaplastic gliomas patients as well as an independent dataset and successfully identified two molecular subgroups with distinct clinical outcome. We found that subgroups have remarkably different survivals with a clear poor prognostic group. Furthermore, the poor prognostic group showed significant benefits for aggressive multimodality therapy, justifying the use of intensive therapy based on molecular stratification.

## Introduction

Gliomas are the most common malignant brain tumor, accounting for 80% of central nervous system (CNS) tumors [1]. World Health Organization (WHO) grade III-anaplastic gliomas are considered high-grade along with WHO Grade IV-glioblastomas, yet unlike glioblastomas, anaplastic gliomas possess widely disparate outcomes with no clear consensus on optimal therapeutic approach. Among anaplastic gliomas, astrocytoma histology is associated with poor outcomes, with a median survival of approximately 2 years and often treated similar to glioblastoma with concurrent temozolomide and radiation therapy (RT) [2]. Anaplastic glioma patients with oligodendroglioma histology and 1p19q co-deletion have a very favorable survival outcome of nearly 15 years when treated with combined radiation and PCV chemotherapy [3, 4]. However, these recommendations represent only a portion of anaplastic gliomas, and there are phase III trials and national guidelines providing evidence for other therapeutic approaches, such as RT or chemotherapy alone [3, 5]. Clinical and molecular prognostic factors, such as histology, age, performance status, *IDH1* mutation status, and 1p19q co-deletion are being used to help guide selection of therapy. However, further objective measures to accurately prognosticate patients with anaplastic gliomas may improve selection of appropriate therapy. Gene expression profiling is being widely investigated for objective tumor classification and prognostication[6] and for gliomas there is evidence that molecular subtyping may serve as a better prognostic marker than histology alone [7–9].

The Cancer Genome Atlas (TCGA) is a large-scale dataset with clinical and genomic data of over 30 cancer types. Grade IV gliomas or glioblastomas were the first cancer type to be studied by TCGA, and investigation of gene expression patterns in glioblastoma suggested 4 distinct subtypes that may be associated with differential therapeutic outcomes [10]. Unlike the considerable progress made in the molecular classification of glioblastoma, similar data are lacking for anaplastic gliomas. Recently, TCGA has added the “Brain Lower Grade Glioma” dataset which includes over 500 patients, and includes patients with grade 3 anaplastic gliomas [11]. The Repository for Molecular Brain Neoplasia Data (REMBRANDT) is another large genomic dataset which integrates clinical and genomic material for all grades and histologies of gliomas [12]. Between the two, they represent two of the largest datasets with genomic and clinical data available for anaplastic gliomas.

Previous studies, utilizing unsupervised cluster analysis of high grade gliomas, have identified subsets with variable prognoses, however, these studies largely are defined by patients with grade 2 and 4 gliomas [7, 11, 13, 14]. In our study, we analyzed expression profiles of only anaplastic gliomas from TCGA. In order to simplify analysis and improve potential clinical translatability, we utilized a previously identified low grade glioma specific-pathway signature containing 39 genes, which was derived by a novel network-based integrative algorithm utilizing on somatic mutation frequency, pathway databases and molecular interaction networks [15]. Based on the expression patterns of these low grade glioma signature genes, we classify anaplastic gliomas into distinct molecular subgroups and compare their survival outcomes. These results were then validated with the REMBRANDT dataset.

## Methods

We previously generated a glioma-specific pathway signature which consists of 39 genes commonly mutated and/or linked via gene-gene interactions [15]. By integrating gene-gene interaction networks and molecular pathway databases, identification of not only frequently mutated genes but their interacting partners that may play a role in low grade glioma biology can be identified. We previously successfully demonstrated that the cancer-type-specific pathway signatures identified by our method could serve as robust prognostic biomarkers that stratify patients with distinct clinical outcomes across many cancer types [15]. The list of all 39 genes and their pathway networks for the low grade glioma specific pathway are shown in Supplementary Figure 1.

The TCGA Brain Lower Grade Glioma Data Matrix and cBioPortal for Cancer Genomics [16, 17] were reviewed to identify patients with anaplastic gliomas. The clinical dataset was further screened to include only patients for whom both survival and censoring data were available. A total of 176 patients met the inclusion criteria. Level 3 *RNASeqV2* data restricted to those patient ID’s were retrieved for cluster analysis. Clinical factors obtained from the TCGA clinical dataset included age, KPS, histology, RT and chemotherapy use, and *IDH1/2* mutation status. 1p19q co-deletion status was determined by an arm log2 fold change of at least -0.2 in both 1p and 19q within each patient’s single nucleotide polymorphism array based data (NoCNV, HG19). The clinical outcome of interest was overall survival (OS).

Unsupervised consensus clustering, based on expression of the 39 gene signature, was performed with cluster sizes between 2 to 8. The expression values were standardized by z-score transformation, where the transformation is applied to each gene individually. Then, for the given cluster size, the k-means algorithm was performed with 500 iterations on 80% of the samples with the fixed cluster size, k. Clustering results from all 500 iterations were combined by constructing a consensus matrix where each element ranges from 0 to 1 and represents the probability that two corresponding samples belong to the same cluster. The final consensus clusters were obtained by applying the hierarchical clustering method to the consensus matrix. The optimal cluster size was chosen by the Cophenetic correlation coefficient and visual inspection of the consensus matrices. An independent validation cohort of patients with anaplastic gliomas from the REMBRANDT dataset was similarly analyzed, once patients with clinical and cDNA microarray data were identified (*Affymetrix HG-U133_Plus_2*).

### Statistical Analysis

Statistical analysis was performed with commercially available software (JMP version 11.2, SAS Institute Inc.). The optimally chosen clusters for the TCGA dataset were assessed for survival by the Kaplan-Meier method and the log-rank test. Differences in clinical and demographic characteristics between molecular subgroups identified by clustering were evaluated by the Chi-square test for categorical variables. Continuous variables such as age and KPS were dichotomized into binary variables based on their median values. In univariate analysis, associations between OS and each of the molecular, clinical and demographical factors were assessed by the Cox proportional hazards model and/or the log-rank test, with p≤.05 considered significant. Multivariable analysis of the effects of molecular, clinical and demographic factors on OS was measured by adjusted hazard ratios in the Cox proportional hazards model, with p≤.05 considered significant. The REMBRANDT dataset was chosen as a validation cohort to determine whether the 39 gene signature can reproducibly define prognostic subgroups of patients with anaplastic gliomas. The overall survival of the chosen clusters from the REMBRANDT dataset was similarly measured by the Kaplan-Meier method and the log-rank test. All reported P-values in the statistical analysis are two-sided. They were not adjusted for multiple comparisons.

For evaluation of the predictive power of the molecular subgroup type with other prognostic factors, we calculated Somers’ D for the following four Cox models with different sets of covariates. Model 1 was performed with molecular subgroup type as a single covariate, Model 2 was performed with *IDH1/2* mutation and 1p19q co-deletion, Model 3 was performed with molecular subgroup type, *IDH1/2* mutation and 1p19q co-deletion, and Model 4 combined the molecular subgroup type, *IDH1/2* mutation, 1p19q co-deletion plus all clinical and demographical factors used in the MVA as covariates.

## Results

### TCGA Dataset

A total of 176 patients with grade III gliomas were identified within the TCGA Brain Lower Grade Glioma dataset. Clinical and demographic characteristics of the patient cohort are shown in Table 1. The median follow-up of the TCGA cohort was 17.4 months (range 0-211 months) and the median survival of the TCGA cohort was 50.8 months with 51 patients dead and 125 patients censored. Unsupervised k-means clustering of the TCGA cohort with cluster sizes 2 through 8 found k=2 to be optimal based on visual inspection and the cophenetic correlation coefficient (Supplementary Figure2). Of the 39 signature genes, a total of 27 were significantly differentially expressed between the two clusters. The five most over-expressed genes were CHEK2, CCNB1, BRCA2, AURKA, and PCNA. The five most under-expressed genes were YAF2, CHD3, PPA1, NAT6, and EEF2 (Figure 1).

**Figure 1.**
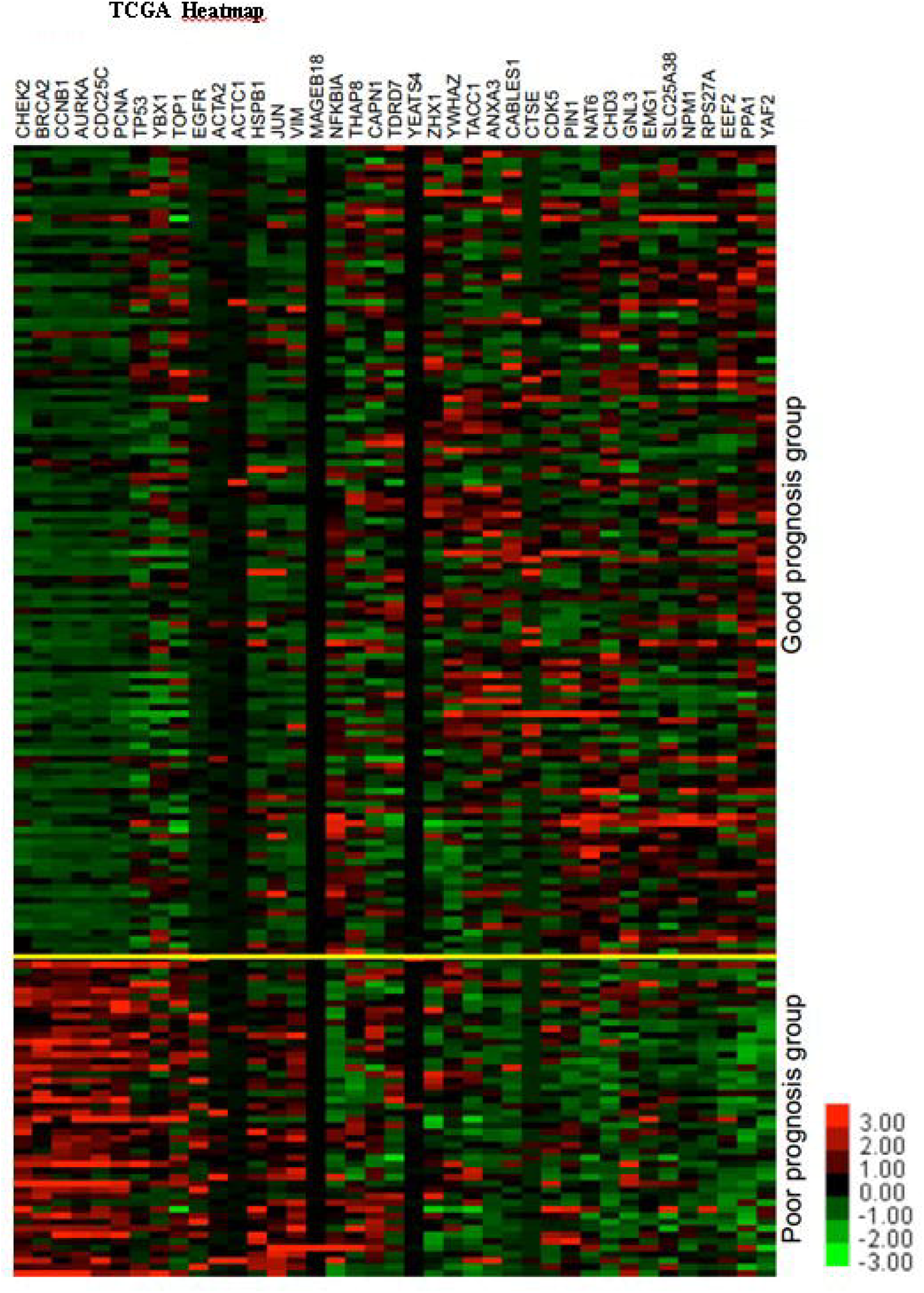
TCGA Heatmap. Columns and rows represent 39 gene-glioma signature and individual patients, respectively. Red and green colors indicate genes overexpressed and underexpressed, respectively.

When OS of the two clusters was evaluated, there was a clear distinction in the two groups with a substantial difference in survival. The median survival of one subgroup was 19.9 months (n=50), which was labeled the poor prognostic subgroup, and the median survival of the second subgroup was 67.4 months (n=126), which was labeled the good prognostic subgroup (p<0.001) (Figure 2A). Clinical and demographic comparisons between the subgroups are shown in Table 2.

**Figure 2.**
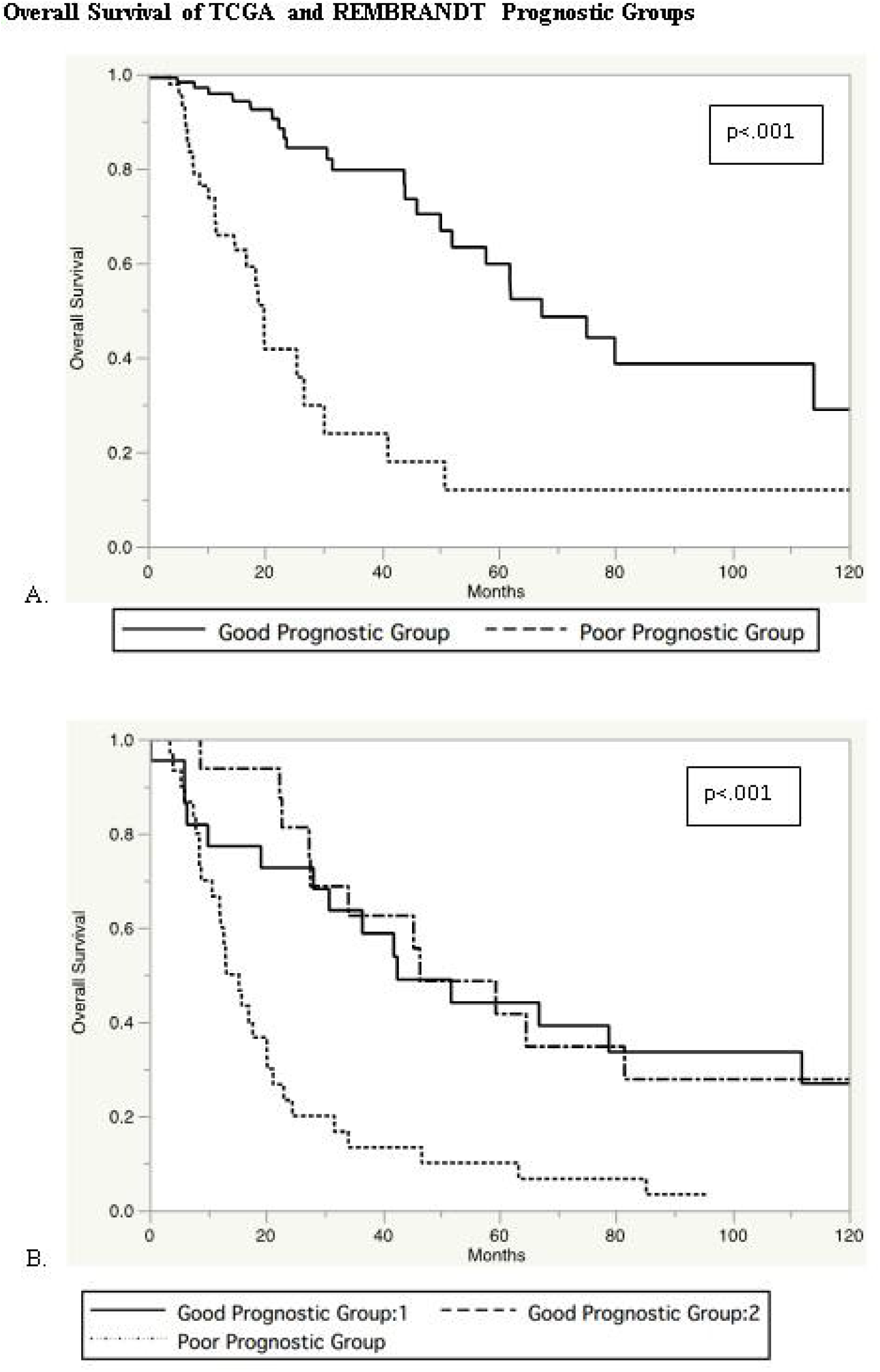
Overall Survival of TCGA and REMBRANDT Prognostic Groups. A. TCGA Prognostic groups: poor prognostic group n=50, good prognostic group n=126 B. REMBRANDT Prognostic groups: poor prognostic group n=30, good prognostic group 1 n=22, good prognostic group 2 n=16

In univariate Cox regression analysis, the molecular subgroup type, age >45, KPS ≥90, *IDH1/2* mutation status, 1p19q co-deletion, chemotherapy, and use of both chemotherapy and radiation therapy were significantly associated with survival, while histology and RT use were not. In multivariable Cox regression analysis, the molecular subgroup type remained significant, along with age >45, KPS ≥90, 1p19q co-deletion, and RT use. Unadjusted and adjusted hazard ratios for each factor along with the corresponding 95% confidence intervals and p-values are summarized in Table 3. Within the poor prognostic subgroup, those patients who received chemotherapy had a significantly prolonged survival of 19.9 versus 11.5 months (p=.03) and those patients who had received both chemotherapy and radiation therapy as part of their treatment course had an improved overall survival of 19.9 versus 15.0 months (p=.05).

### Predictive Power Measured by Somers’ D

The molecular subgroup types identified via clustering analysis was evaluated in Model 1, with a Somers’ D value of 0.41. *IDH1/2* mutation and 1p19q co-deletion status were evaluated in Model 2, with a Somers’ D value of 0.45, showing comparable power to predict overall survival between our molecular subtyping and known molecular prognostic factors. When combining our molecular subtyping with *IDH1/2* mutation and 1p19q co-deletion into Model 3, there was further power to predict for survival with a value of 0.52. Model 4 combined the molecular subgroup types, *IDH1/2* mutation status, and 1p19q co-deletion, along with all clinical and demographical factors used in the MVA, and they provided the highest predictive power of 0.77.

### Validation Dataset-REMBRANDT

The REMBRANDT dataset was chosen as a validation cohort to determine whether the 39 gene signature can reproducibly define prognostic subgroups of patients with anaplastic gliomas. A total of 68 patients with grade III gliomas were identified within the REMBRANDT database. The median survival within this cohort was 27.4 months and the most common histology was astrocytoma (64%). Unsupervised k-means clustering with cluster sizes of 2 through 8 found k=3 to optimal based on visual inspection and the cophenetic correlation coefficient (Supplementary Figure 3). Of the three clusters, one had a significantly inferior survival with a median survival of 14.3 months (n=30) and the other two clusters had a similar median survival of 42.5 (n=22) months and 46.4 months (n=16), with a significant difference in overall survival between the three clusters based on the log-rank test (p<.001) [Figure 2B]. Gene expression patterns identified 25 of 39 signature genes which were significantly over or underexpressed in the poor prognostic group (t-test with FDR adjusted p < 0.05). Between the TCGA and REMBRANDT databases, there was concordance of expression in 21/39 genes (Figure 3). Fourteen of the genes were overexpressed in both TCGA and REMBRANDT datasets and 7 were under-expressed in both datasets.

**Figure 3.**
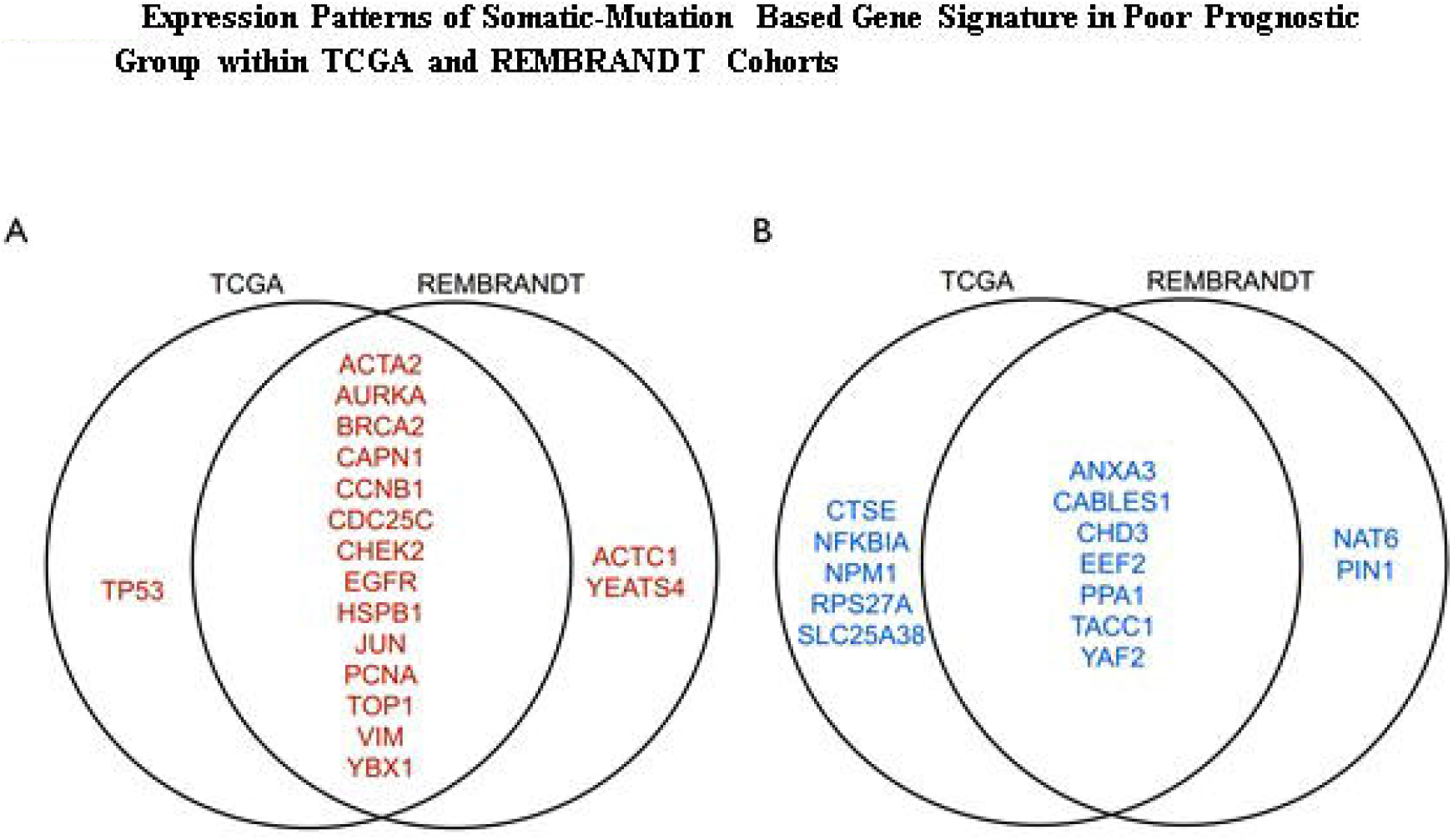
Expression Patterns of Somatic-Mutation Based Biomarkers in Poor Prognostic Group within TCGA and REMBRANDT Cohorts. (A) Venn diagram shows genes that are commonly or differently overexpressed genes in poor prognosis groups within TCGA and REMBRANDT cohorts (B) Venn diagram shows genes that are commonly or differently underexpressed genes in poor prognosis groups within TCGA and REMBRANDT cohorts

## Discussion

Current therapeutic strategies for anaplastic gliomas vary widely and they may entail concurrent or sequential chemotherapy and RT, chemotherapy or RT alone, or even observation in certain favorable cases. As can be seen, there is no clear consensus on treatment based on known prognostic factors alone. Thus, further stratification may help identify those patients with the worst outcomes for whom aggressive combined modality therapy may be necessary and to potentially avoid unnecessary treatment toxicity in others.

We previously developed a novel algorithm, named NTriPath (Network regularized non-negative TRI matrix factorization for PATHway identification), which integrates somatic mutation, gene-gene interaction networks, and pathway databases to identify altered pathways in 19 different cancer types from TCGA database [15]. In doing so, we identified 39 distinct genes for low grade gliomas which include not only genes with mutations occurring at high frequency but also non-mutated genes linked to the mutations which may have fundamental roles in the biology of gliomas. By utilizing this glioma-specific 39 gene signature, our goal was to determine whether there were unique subgroups of anaplastic gliomas with differential survival outcomes.

With the 39-gene signature, it was possible to stratify anaplastic gliomas, without any clinical bias, into two distinct subtypes with significantly different survivals. These results were significant and independent of other established prognostic factors, such as histology, *IDH1/2* mutation, and 1p19q co-deletion. Presence of *IDH1/2* mutations and 1p19q co-deletions is commonly used to identify favorable patients, but in our good prognostic subgroup, these alterations were absent in 33% and 39% respectively, suggesting that these known prognostic factors do not fully define a favorable prognosis. Based on the Somers’ D measure, it was shown that the molecular subgroup type had a similar predictive power as *IDH1/2* mutation status and 1p19q co-deletion, and when combined, there was an added benefit in predicting survival. Amongst the patients in the poor prognostic group, those who received chemotherapy and radiation therapy had an improved survival, justifying therapy intensification within this molecular subtype defined by our gene signature. The poor prognosis subgroup was characterized by unfavorable factors, such as the lack of IDH1/2 mutation and 1p19q deletion, as well as older median age and worse KPS. However, even after taking these prognostic factors into consideration, the molecular subgroup types remained a significant predictor of survival on multivariate analysis and when combined with other known prognostic factors, it further improved the predictive power.

Between the primary dataset (TCGA) and the validation dataset (REMBRANDT), the 39 gene signature had concordance of expression patterns in 21 genes. Many of the overexpressed genes are implicated in cellular proliferation, cell-cycle regulation, and DNA damage repair. For example, *AURKA* codes for Aurora A kinase, a key protein involved with cell-cycle regulation, mitosis, and proliferative signal pathways [18, 19]. Aurora A kinase expression has been found to increase with increasing glioma grade, with increased expression correlating with worse overall survival in high-grade gliomas[20]. Alisertib is an Aurora-A kinase specific inhibitor which has shown inhibition proliferation in glioblastoma neurospheres and is currently being testing in a phase I trial in patients with recurrent high grade gliomas. [21] *Cyclin B1* is involved in the transition from the G2 to M phase by complexing with Cdc2 and in gliomas, higher grade tumors have been found to have higher expression of cyclin B1 [22]. In multiple tumor cell lines, small interfering RNAs were able to decrease proliferation and arrest cells in the G2/M phase. Targeting this pathway in the cell cycle in patients found to have a poor-prognosis could potentially provide efficacy against the primary tumor as well as work synergistically with RT, given the latter’s increased efficacy in the G2/M phase of the cell cycle. Epidermal growth-factor receptor (*EGFR*) is one of the most well-known drug targets and in glioblastoma *EGFR* amplification occurs in approximately 40% of tumors [23]. When looking at the poor prognostic subgroup of both TCGA and REMBRANDT anaplastic gliomas, *EGFR* overexpression was consistently characteristic in both datasets. The rate of *EGFR* overexpression in the TCGA data set was 19% and patients with *EGFR* overexpression in this database had a significantly poorer overall survival, with a median survival of 24 versus 62 months (p=0.003), representing another potential therapeutic target in anaplastic gliomas.

The 21 concordant genes between both datasets represent genes which in the future may be validated with clinical samples, reducing the number of genes to allow widespread and efficient utilization of these markers much like Oncoytpe DX testing in patients with breast cancer [24].As an example of the prospective potential for glioma subtyping, the randomized EORTC 26951 successfully performed molecular subtyping of fresh-frozen samples from a prospective clinical trial, validating the ability and addition of molecular subtyping to known prognostic factors [8]. Furthermore, within this study, 37% of patients without 1p19q co-deletion within a distinct subtype benefitted from the addition of PCV chemotherapy. With current standard of care limiting PCV plus radiation therapy to only those patients with 1p19q co-deletion, there is the possibility that a considerable number of patients who may benefit are not offered the optimal treatment which would only be considered with the addition of molecular based subtyping.

One important caveats of this study is incompleteness of clinical information in the databases, such as surgical extent in TCGA and lack of clinical data within REMBRANDT, which were not accounted for in our analysis. Nevertheless, molecular subtyping according to the expression of our 39 gene signature identified distinct prognostic groups of anaplastic gliomas. For those patients in the poor prognostic group, aggressive multimodality therapy, similar to GBM may be justified. The overexpressed genes and pathways that characterize the poor prognostic group may be important targets for novel agents not previously studied in anaplastic gliomas.

## Acknowledgements

The results shown here are in whole or part based upon data generated by the TCGA Research Network: http://cancergenome.nih.gov/.

Design and conduct of the study; collection, management, analysis, and interpretation of the data; and preparation, review, or approval of the manuscript was done without any financial or material support.

---

**Supplementary Figure 1.**
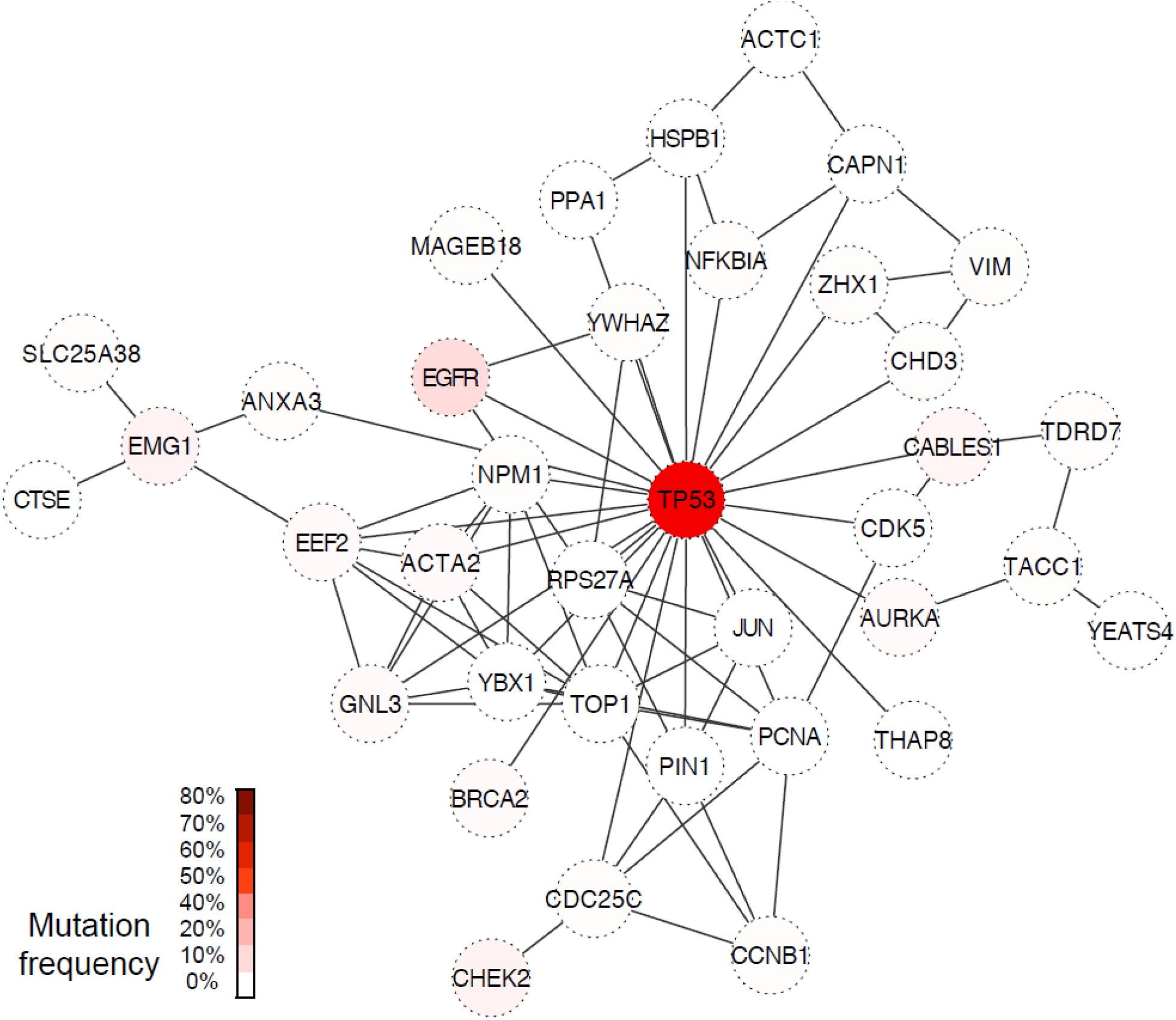
NTriPath Glioma Specific Biomarkers. 39 genes identified via NTriPath linked via somatic mutations and pathway networks which are implicated in glioma biology. Red color indicates mutation frequency in TCGA LGG dataset.

**Supplementary Figure 2.**
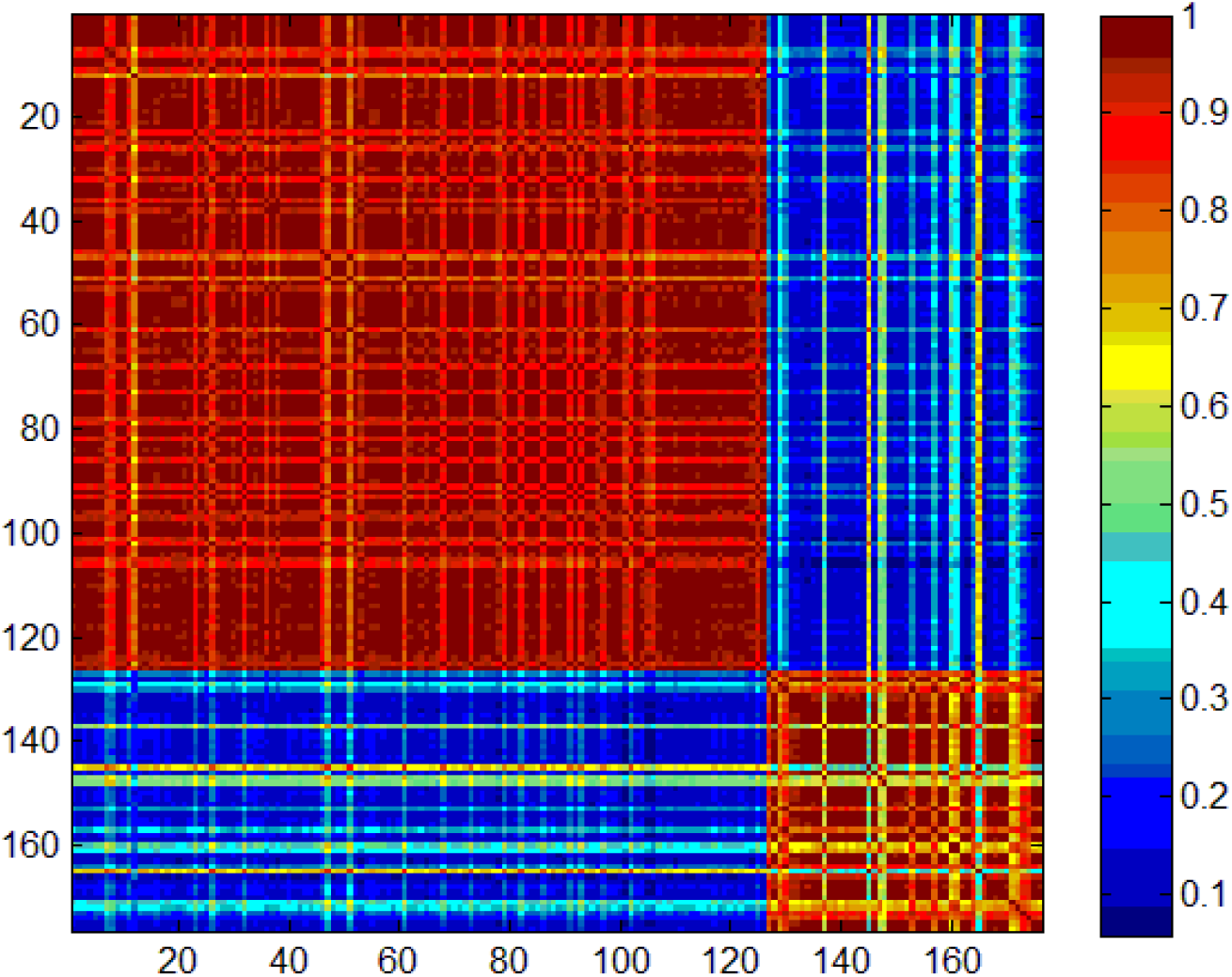
Sample by Sample Data Matrix for TCGA cohort (k=2) A heatmap represents a consensus matrix obtained by consensus clustering. Red color indicates increasing probability of patients belonging to same cluster. Blue color indicates decreasing probability of patients belonging to same cluster.

**Supplementary Figure 3.**
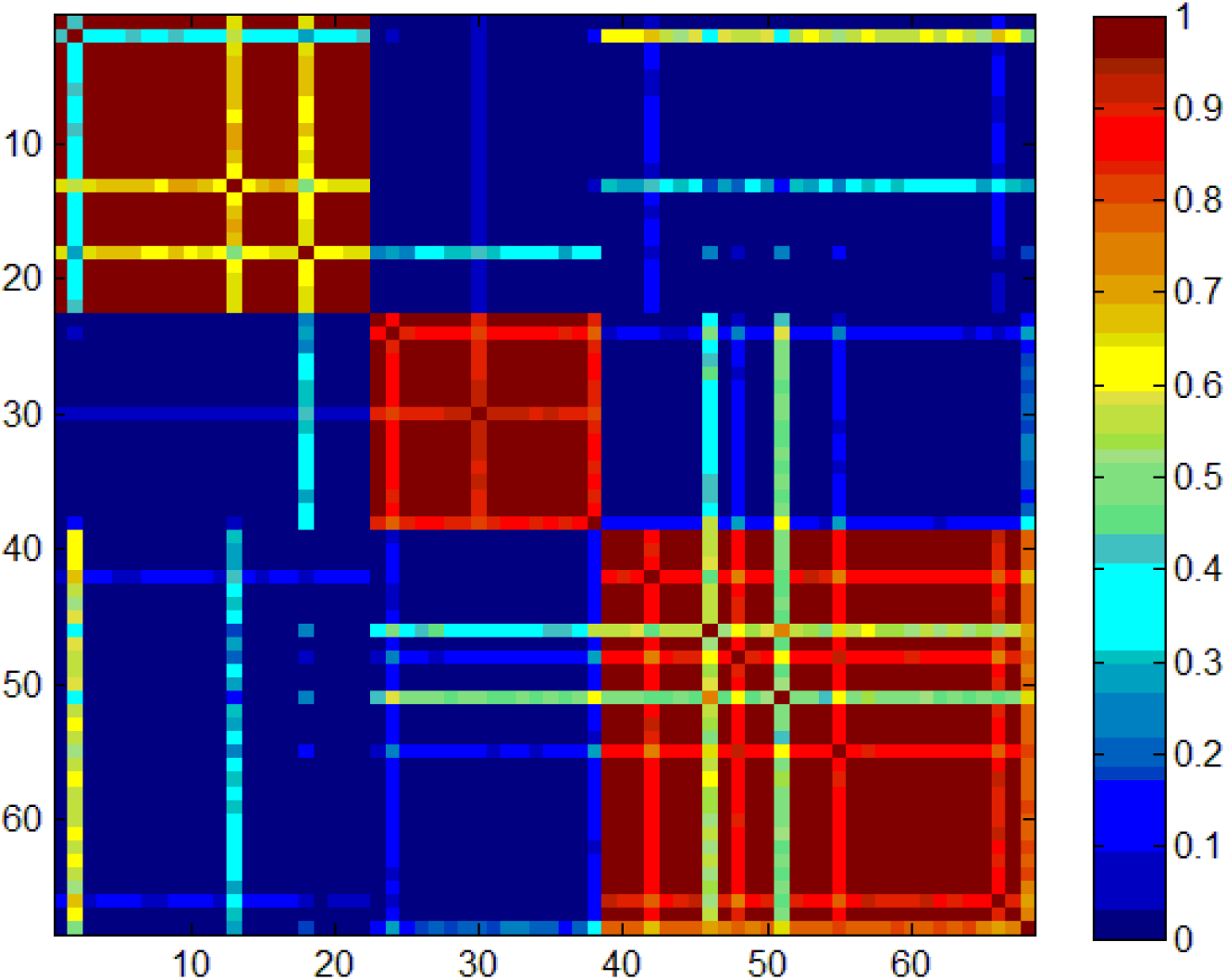
Sample by Sample Data Matrix for REMBRANDT cohort (k=3) A heatmap represents a consensus matrix obtained by consensus clustering. Red color indicates increasing probability of patients belonging to same cluster. Blue color indicates decreasing probability of patients belonging to same cluster.

